# Cortical microtubule pulling forces contribute to the union of the parental genomes in the *C. elegans* zygote

**DOI:** 10.1101/2021.11.19.469242

**Authors:** G. Velez-Aguilera, B. Ossareh-Nazari, L. Van Hove, N. Joly, L. Pintard

## Abstract

Previously, we reported that the Polo-like kinase PLK-1 phosphorylates the single *C. elegans* lamin (LMN-1) to trigger lamina depolymerization during mitosis. We showed that this event is required for the formation of a pronuclear envelopes scission event that removes membranes on the juxtaposed oocyte and sperm pronuclear envelopes in the zygote, allowing the parental chromosomes to merge in a single nucleus after segregation (Velez-Aguilera, 2020). Here we show that cortical microtubule pulling forces contribute to pronuclear envelopes scission by promoting mitotic spindle elongation. We also demonstrate that weakening of the pronuclear envelopes, via PLK-1-mediated lamina depolymerization, is a prerequisite for the astral microtubule pulling forces to trigger pronuclear membranes scission. Finally, we provide evidence that PLK-1 mainly acts via lamina depolymerization in this process. These observations thus indicate that temporal coordination between lamina depolymerization and mitotic spindle elongation facilitates pronuclear envelopes scission and parental genomes unification.

## Introduction

The life of sexually reproducing organisms starts with the joining of two haploid genomes. Parental chromosomes are first replicated in distinct pronuclei, each surrounded by a nuclear envelope, and meet for the first time during the first mitosis. Coordinated disassembly of the pronuclear envelopes is required to promote the reunification of the parental chromosomes in the fertilized zygote, but the underlying mechanisms are incompletely understood.

The *C. elegans* zygote provides an attractive model system to investigate the mechanisms by which the maternal and paternal genomes unify at the beginning of life (Oegema and Hyman, 2006; Cohen-Fix and Askjaer, 2017; Pintard and Bowerman, 2019). After fertilization, the oocyte and sperm chromosomes surrounded by a nuclear envelope localize to opposite sides of the zygote. The female pronucleus is located in the anterior, whereas the male pronucleus - and associated centrosomes - is in the posterior. When the two pronuclei meet, they change their shape, flattening out along their juxtaposed sides, eventually aligning along the AP axis of the one-cell embryo (Oegema and Hyman, 2006). The two pronuclei then undergo spatially regulated nuclear envelope breakdown to permit chromosome attachment to microtubules and their alignment on the metaphase plate: breakdown of the pronuclear envelopes starts in the vicinity of the centrosomes and then later between parental chromosomes (Lee et al., 2000; Hachet et al., 2012; Velez-Aguilera et al., 2020). The breakdown of the juxtaposed pronuclear envelopes of the oocyte and sperm begins with the formation of a membrane scission event (also called a membrane gap), so that the pronuclear envelopes are removed from the regions between the chromosomes. This event occurs concomitantly with the parental chromosomes congressing on the metaphase plate, 10 to 40s before anaphase onset (Audhya et al., 2007; Rahman et al., 2020), allowing the chromosomes from the two pronuclei to mingle on the metaphase plate and join in a single nucleus after chromosome segregation. How this membrane scission event is formed is still poorly understood, but intriguingly, it always appears at an equal distance from the two centrosomes and depends on the proper alignment of the chromosomes on the metaphase plate (Rahman et al., 2015). Formation of this membrane gap also requires depolymerization of the lamina by the mitotic Polo-like kinase PLK-1 (Rahman et al., 2015; Martino et al., 2017; Velez-Aguilera et al., 2020). Accordingly, expression of an LMN-1 version, carrying eight serine replaced by non-phosphorylable alanines [hereafter LMN-1 8A], is sufficient to prevent the formation of pronuclear envelopes scission, resulting in the typical appearance of embryos with a paired nuclei phenotype (Rahman et al., 2015; Martino et al., 2017; Velez-Aguilera et al., 2020). In these embryos the two sets of parental chromosomes remain physically separated during mitosis and segregate into two separate DNA masses at each pole of the spindle. This in turn leads to the formation of 2 nuclei in each cell of the two-cell embryo (paired nuclei) (Audhya et al., 2007; Bahmanyar et al., 2014; Galy et al., 2008; Rahman et al., 2015; Martino et al., 2017; Velez-Aguilera et al., 2020).

Besides PLK-1-mediated lamina depolymerization, mechanical forces provided by astral microtubules could also contribute to pronuclear envelopes scission, by facilitating the removal of the lamina, similar to the situation in human cells (Salina et al., 2002; Beaudouin et al., 2002), but possibly also by promoting mitotic spindle elongation.

Here, we follow up on our previous work showing that pronuclear envelope scission is regulated via PLK-1-mediated lamina depolymerization (Velez-Aguilera et al., 2020) by testing whether cortical microtubule pulling forces also contribute to this process. We show that astral microtubule pulling forces contribute to pronuclear membranes scission both by facilitating lamina disassembly and by promoting mitotic spindle elongation. Our observations thus suggest that temporal coordination between lamina depolymerization and spindle elongation induces pronuclear envelopes scission, and in turn, facilitates the reunification of the parental chromosomes after segregation.

## Results and discussion

### Microtubules dynamics during nuclear envelope breakdown in the one-cell *C. elegans* embryo

Previous work established that nuclear envelope breakdown (NEBD) is spatially regulated in the fertilized one-cell *C. elegans* zygote, but the exact timing of each event and the contribution of microtubules to this process had not yet been investigated. To address this point, we simultaneously visualized microtubules and nuclear envelope dynamics with a high temporal resolution during NEBD. We used spinning disk confocal microscopy to film one-cell embryos (1 image/2s), expressing fluorescently labeled tubulin (GFP::TBB-2) with either the lamina (mCherry::LMN-1) (**Figure 1A, movie 1**), or the transmembrane nucleoporin NPP-22 (mCherry::NPP-22) (**Figure 1B** **movie 2**). We also filmed embryos expressing GFP::TBB-2 and mCherry-tagged histone (mCherry::Histone) to monitor the configuration of the chromosomes during the different steps of NEBD (**Figure 1C, movie 3**).

**Figure 1:**
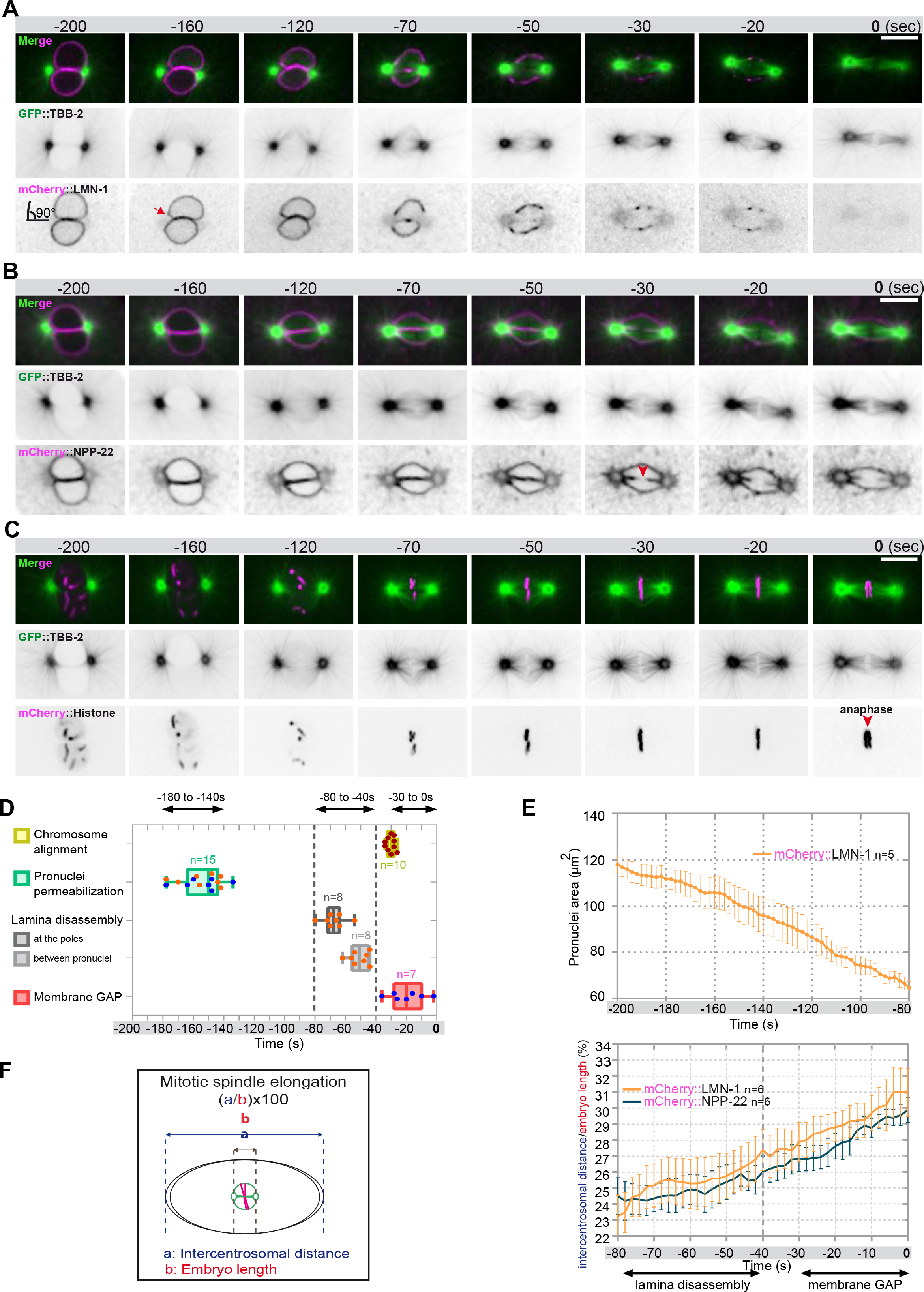
Monitoring microtubule and pronuclear envelopes dynamics in the one-cell embryo. **A-B-C-** Spinning disk confocal micrographs of embryos expressing wild-type GFP::TBB-2 (shown alone, and in green in the merged images) and (**A**) mCherry::LMN-1 (shown alone and in magenta in the merged image), or (**B**) mCherry::NPP-22 (magenta, in the merged image), or (**C**) mCherry::Histone, (magenta, in the merged image). Timings in seconds are relative to anaphase onset (0s). All panels are at the same magnification. Scale Bar, 10 μm. **D-** Steps of pronuclear envelopes breakdown: pronuclei permeabilization, lamina disassembly (at the poles, between pronuclei) and pronuclear envelopes scission event (membrane gap) relative to anaphase onset was scored in multiple embryos (n). The timing of pronuclear envelopes scission was scored in embryos expressing mCherry::NPP-22 (blue dots, n=7) while the timing of lamina disassembly was scored in embryos expressing mCherry::LMN-1 (orange dots, n=8). **E-** Graph presenting the surface area occupied by the pronuclei starting 200s before anaphase onset (0s). **F-** Graph presenting the intercentrosomal distance normalized to embryo length starting 80s before anaphase onset (0s) in embryos expressing GFP::TBB-2 and mCherry::LMN-1 or mCherry::NPP-22.

Before nuclear envelope breakdown, 200s prior to anaphase onset, microtubules were excluded from the pronuclei space, and the pronuclear envelopes appeared flattened along their juxtaposed sides but presented elsewhere a rounded shape, except around centrosomes where the nuclear envelopes were highly curved (90° angle) (**Figure 1A, 1B**). Later, 160s before anaphase onset, signs of nuclear envelope deformation were apparent in the vicinity of the centrosomes (**Figure 1A**, red arrow). At these sites, the pronuclear envelopes appeared deformed toward the centrosomes, suggesting that microtubules, emanating from the centrosomes were pulling or pushing on the pronuclear envelope. Pronuclear envelopes permeabilization, measured by the appearance of soluble tubulin in the pronuclei space, was detected around 160s before anaphase onset, concomitantly with pronuclear membranes deformation near the centrosomes (**Figure 1A to 1D, Table 1 Source Data 1**). Nuclear envelope permeabilization was systematically detected first in the male pronucleus (**Figure S1**). Soon after membrane permeabilization, the first microtubules started to invade the pronuclei space. As more microtubules invaded the pronuclei space to capture paternal chromosomes, the overall surface of the pronuclei decreased (**Figure 1E**, **Table 1 Source Data 1**) and the pronuclei adopted a more triangular shape, dictated by microtubules assembling the mitotic spindle (**Figure 1B**).

Signs of lamina disappearance from the pronuclear envelopes systematically started in the vicinity of centrosomes, 70s before anaphase onset (**Figure 1A, 1D and S2**), possibly as a consequence of microtubule pulling or pushing at this site, together with the action of the lamin kinase PLK-1, which is enriched at centrosomes (Chase et al., 2000; Budirahardja and Gonczy, 2008; Nishi et al., 2008; Martino et al., 2017). The lamina progressively disappeared in the vicinity of the centrosomes but persisted between the parental chromosomes. Lamina disassembly between the pronuclei started only 50 s before anaphase onset (**Figure 1A and 1D**). At this time point, the lamina network was still detected on the envelopes surrounding the chromosomes (**Figure 1A**). The membrane scission event between the juxtaposed pronuclei was detected later, around 30s before anaphase onset (**Figure 1B, 1D and S3**), consistent with previous observations (Audhya et al., 2007). The scission event appeared immediately after chromosome alignment on the metaphase plate (**Figure 1B red arrowhead, 1C-D, S3, Table 1 Source Data 1**).

Starting 80s before anaphase onset, the distance between the centrosomes steadily increased as a result of spindle elongation (**Figure 1F, Table 1 Source Data 1**). The beginning of spindle elongation was concomitant with lamina depolymerization but preceded membrane gap formation. Based on these observations, we hypothesized that by pulling or pushing on nuclear envelopes and membranes, and by tearing apart the lamina and by elongating the mitotic spindle, astral microtubule pulling forces might contribute to pronuclear envelopes breakdown, and thus to the reunification of the parental chromosomes in the early *C. elegans* embryo.

### Paired nuclei phenotype upon reduction of cortical microtubule pulling forces

The paired nuclei phenotype is a visual readout of the failure to properly remove the nuclear envelope from the pronuclei (Audhya et al., 2007; Bahmanyar et al., 2014; Galy et al., 2008; Rahman et al., 2015; Martino et al., 2017; Velez-Aguilera et al., 2020).

To test whether microtubule pulling forces contribute to pronuclear envelopes breakdown, we examined whether experimental reduction of microtubule-dependent cortical pulling forces could modify the penetrance of the paired nuclei phenotype of embryos expressing the *gfp::lmn-1* 8A allele (Link et al., 2018; Velez-Aguilera et al., 2020). Because this allele partially stabilizes the lamina, 8% of the embryos expressing GFP::LMN-1 8A present double paired nuclei at the 2-cell stage, and another 9% present a single paired nuclei cell (**Figure 2A, Table 2 Source Data 2**). To reduce the cortical pulling forces, we used RNAi to partially deplete GPR-1/2, which are part of an evolutionarily conserved complex anchoring the dynein motor to the embryo cortex (Colombo et al., 2003; Gotta et al., 2003; Srinivasan et al., 2003) (**Figure 2B**). Mild depletion of GPR-1/2 in *gfp::lmn-1* 8A embryos greatly enhanced the percentage of embryos presenting a paired nuclei phenotype, with nearly 54% showing double paired nuclei and 24% a single paired nuclei (**Figure 2A, Table 2 Source Data 2**). More severe RNAi-mediated *gpr-1/2* inactivation (material and methods) further increased the percentage of *gfp::lmn-1* 8A mutant embryos presenting a double paired nuclei phenotype to 83% (**Figure 2A, Table 2 Source Data 2**). Similar treatments of the embryos expressing a wild-type GFP::LMN-1 allele had only little effect with 5 % of embryos presenting a double paired nuclei phenotype and 4% presenting a single paired nuclei phenotype (**Figure 2A**). Thus, reduction of cortical microtubule pulling forces enhance the paired nuclei phenotype of embryos with a partially stabilized lamina network. These observations indicate that microtubule pulling forces facilitate the union of the parental chromosomes in the one-cell embryo.

**Figure 2:**
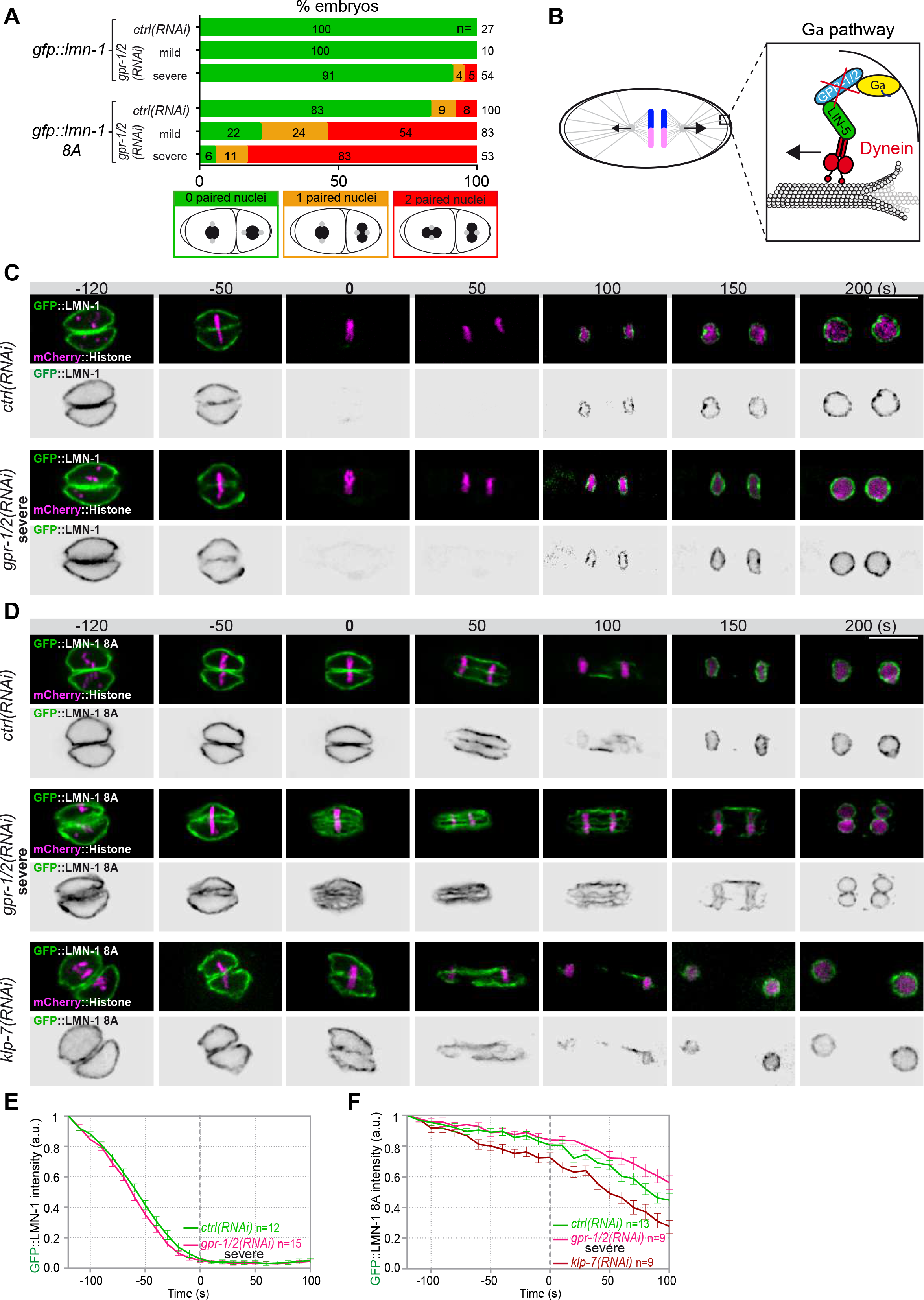
Astral microtubule pulling forces contribute to the union of the parental chromosomes during mitosis. **A-** Percentage of *lmn-1Δ* ; *gfp::lmn-1* and *lmn-1Δ* ; *gfp::lmn-1 8A* embryos presenting 0 (green bars), 1 (orange bars) or 2 (red bars) paired nuclei at the two-cell stage upon exposure to mock RNAi (*ctrl*) or *gpr-1/2(RNAi)* at 20°C. The number of embryos of the indicated phenotype (n) is shown in the graph and was collected from more than three independent experiments. **B-** Schematics of a one-cell *C. elegans* embryo in anaphase. The astral pulling forces mediated by the Gα pathway, is schematized in the inset. This pathway, which comprises a complex of Gα (yellow), GPR-1/2 (blue), and LIN-5 (green), anchors dynein (red) to the cell cortex to generate pulling forces when dynein walks toward microtubule minus ends anchored at the spindle poles. Inactivation of GPR-1/2 (red cross) suppresses the astral pulling forces. **C-D** Spinning disk confocal micrographs of early *lmn-1Δ* mutant embryos expressing **(C)** wild-type GFP::LMN-1 or **(D)** GFP::LMN-1 8A (shown alone, and in green in the merged images) and mCherry::Histone (magenta, in the merged image) exposed to mock RNAi (*ctrl*) in the upper panels or *gpr-1/2* and *klp-7* RNAi in the lower panels. Times are in seconds relative to anaphase onset (0s). All panels are at the same magnification Scale Bar, 10 μm. **E-F** Quantification of (**E**) GFP::LMN-1 or (**F**) GFP::LMN-1 8A signal intensity above background at the nuclear envelope in embryos of the indicated genotype during mitosis. The mean +/- SEM is presented for n embryos of the indicated genotypes. Data were collected from three independent experiments.

To test if microtubule pulling forces were directly facilitating lamina disassembly, we monitored GFP::LMN-1 WT and 8A levels throughout mitosis in control versus *gpr- 1/2 (RNAi)* embryos, upon partial or severe *gpr-1/2* inactivation. *gpr-1/2* inactivation had no discernable effect on GFP::LMN-1 WT disassembly (**Figure 2C, 2E, Table 2 Source Data 2**). Likewise, partial *gpr-1/2* inactivation did not significantly stabilize GFP::LMN-1 8A during mitosis (**Figure S4A and S4B, Table 2 Source Data 2**). However, more severe reduction of cortical microtubule pulling forces, using strong RNAi-mediated *gpr-1/2* inactivation, did stabilize GFP::LMN-1 8A (**Figure 2D, 2F, Table 2 Source Data 2**), indicating that microtubule pulling forces contribute to lamina disassembly during mitosis.

To corroborate these observations, we tested whether excessive microtubule pulling forces would facilitate the removal of GFP::LMN-1 8A. To do so, we inactivated the kinesin-13 family member KLP-7, which results in the assembly of an abnormally high number of astral microtubules and thus increases astral cortical pulling forces (Srayko et al., 2005; Gigant et al., 2017). Loss of *klp-7* caused a premature disassembly of GFP::LMN-1 8A during mitosis (**Figure 2D, 2F, Table 2 Source Data 2**), again arguing that astral microtubule pulling forces contribute to the union of the parental chromosomes, at least in part by pulling at the lamina during mitosis. However, our observation that partial inactivation of *gpr-1/2* enhanced the percentage of GFP::LMN-1 8A embryos presenting a paired nuclei phenotype (**Figure 2A, Table 2 Source Data 2)** without further stabilizing the lamina (**Figure S4A and S4B, Table 2 Source Data 2**) suggested that cortical microtubule pulling forces might prevent the formation of embryo with a paired nuclei phenotype by additional mechanism(s).

### Reducing or increasing cortical microtubule pulling forces affects the timing of membrane scission between the parental pronuclei

Scission of the pronuclear envelopes occurs right before anaphase, after the beginning of mitotic spindle elongation (**Figure 1D, 1F**). We thus reasoned that by pulling on centrosomes and membranes, and by elongating the mitotic spindle, microtubule pulling forces might mechanically facilitate membrane scission between the parental pronuclei. To investigate this possibility, we used spinning disk confocal microscopy to monitor pronuclear membranes scission upon strong *gpr-1/2* inactivation in embryos expressing the inner nuclear membrane protein GFP::LEM-2 and mCherry::Histone, allowing simultaneous visualization of pronuclear envelopes and chromosomes (**Figure 3A**). In wild-type embryos, the membrane scission event between the pronuclei was systematically observed around 30 sec before anaphase onset, after chromosomes alignment on the metaphase plate (n=32) (**Figure 3B**, red arrowhead). However, in a vast majority of *gpr-1/2(RNAi)* embryos, the membrane scission between the pronuclei was never detected. Of the 30 embryos analyzed, only 4 presented a membrane scission event (17%) (**Figure 3B, 3C, Table 3 Source Data 3**) and these 4 embryos were also the least affected in spindle elongation (**Figure 3D, Table 3 Source Data 3**), suggesting that spindle elongation contributes to pronuclei membrane scission.

**Figure 3:**
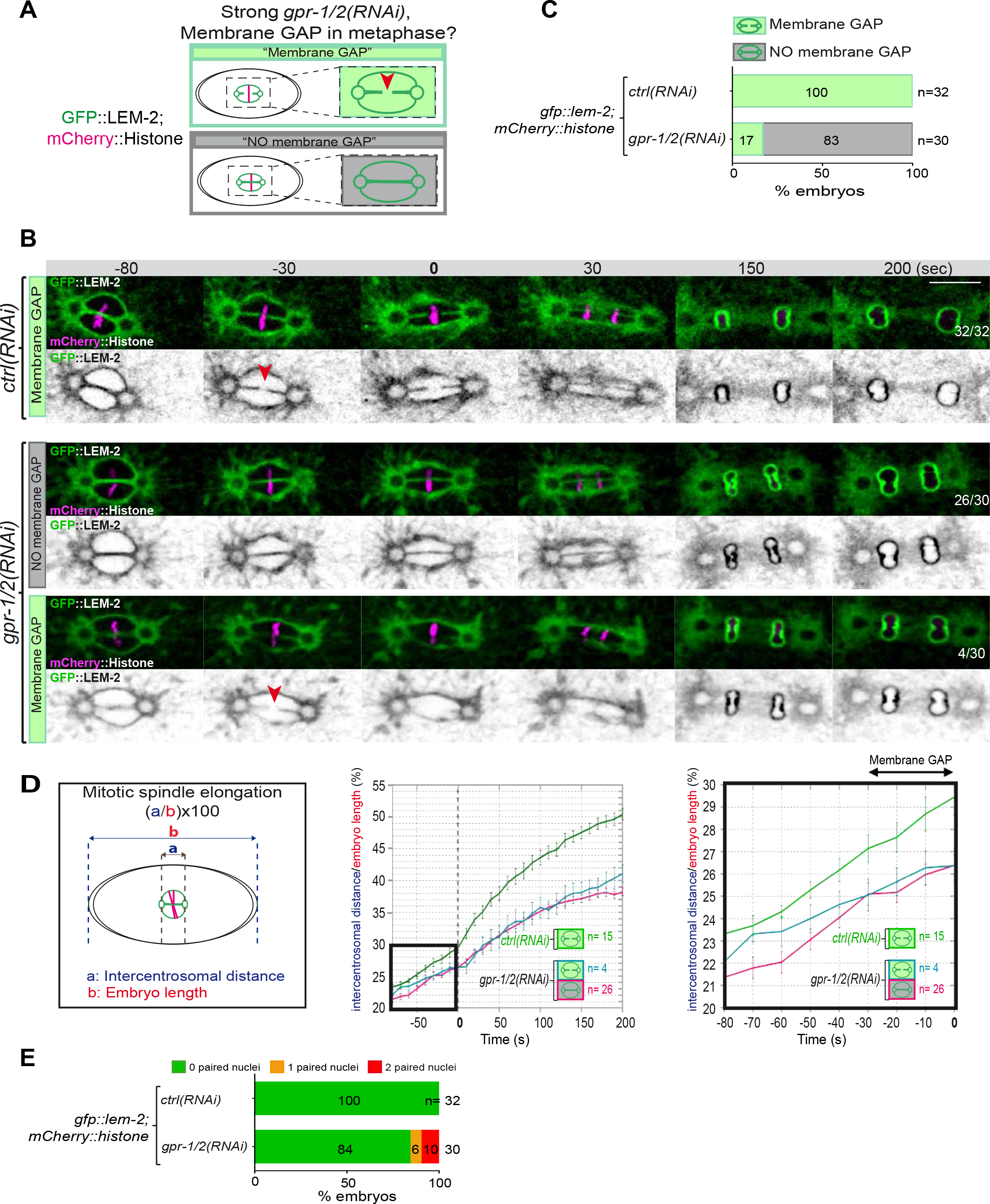
Reduction of astral pulling forces prevents pronuclear membranes scission. **A-** Schematics of the approach used to test the effect of a reduction of astral microtubule pulling forces on pronuclear envelopes scission (membrane gap) during mitosis. **B-** Spinning disk confocal micrographs of one-cell stage embryos expressing the inner nuclear membrane protein GFP::LEM-2 (shown alone, and in green in the merged images) and mCherry::Histone (magenta, in the merged image) exposed to mock RNAi (*ctrl*) in the upper panels or *gpr-1/2(RNAi)* in the lower panels. Times are in seconds relative to anaphase onset (0s). The fraction of embryos that showed the presented phenotype is indicated at the bottom right of each image. All panels are at the same magnification Scale Bar, 10 μm. The orange arrowheads point to the pronuclear envelopes scission event. **C-** Percentage of embryos presenting a pronuclear envelopes scission event upon exposure to mock RNAi (*ctrl*) or *gpr-1/2(RNAi)*. **D-** Graphs showing the intercentrosomal distance normalized to embryo length in percentage during mitosis upon exposure to mock RNAi (*ctrl*) or *gpr-1/2(RNAi)*. Times are in seconds relative to anaphase onset (0s). **E-** Percentage of embryos presenting 0 (green bars), 1 (orange bars) or 2 (red bars) paired nuclei at the two-cell stage upon exposure to mock RNAi (*ctrl*) or *gpr- 1/2(RNAi).* The number of embryos analyzed (n) is indicated on the graph and was collected from three independent experiments.

Notably, a defect in the formation of a membrane scission between the pronuclei was not systematically accompanied by the formation of embryos with a paired nuclei phenotype. Nevertheless, all reforming nuclei were severely misshapen in *gpr- 1/2(RNAi)* embryos and a fraction of them presented a double paired nuclei phenotype at the 2-cell stage (**Figure 3E, Table 3 Source Data 3**). It is possible that during nuclear envelope reformation, the physical barrier separating the parental chromosomes is eventually dismantled, which may explain why single pronuclei manage to form in each blastomere at the 2-cell stage.

If cortical microtubule pulling forces contribute to the formation of the pronuclei membrane scission event by elongating the mitotic spindle, excessive pulling forces might induce a premature membrane scission relative to anaphase onset. To test this hypothesis, we used two complementary approaches to increase astral microtubule pulling forces. We inactivated *klp-7* as before, or *efa-6* (Exchange factor for Arf), which encodes a cortically-localized protein that limits the growth of microtubules near the cell cortex of early embryonic cells. Loss of EFA-6 causes excess centrosome separation and displacement towards the cell cortex early in mitosis and subsequently increased rates of spindle elongation (O’Rourke et al., 2010). We inactivated *klp-7* or *efa-6* in embryos expressing GFP::LEM-2 and mCherry::Histone to monitor pronuclear membranes scission relative to anaphase onset and mitotic spindle length (**Figure 4A**). While the wild-type embryos underwent membrane scission between the juxtaposed pronuclei 0 to 30 sec before anaphase onset, this event occurred systematically earlier in *efa-6* and *klp-7(RNAi)* embryos. In 50% of these embryos, pronuclear membranes scission occurred between 40 and 120 sec before anaphase onset (**Figure 4B, 4C, Table 4 Source Data 4**). Membrane scission occurred earlier in these embryos as a consequence of premature mitotic spindle elongation (**Figure 4D, Table 4 Source Data 4**). By measuring mitotic spindle length at the time of membrane gap formation, we noticed that membrane scission systematically occurred at a similar mitotic spindle length (**Figure 4E, Table 4 Source Data 4**). Taken together, these observations indicate that microtubule pulling forces, by promoting mitotic spindle elongation, contribute to pronuclear membranes scission, possibly by tearing apart the pronuclear membranes.

**Figure 4:**
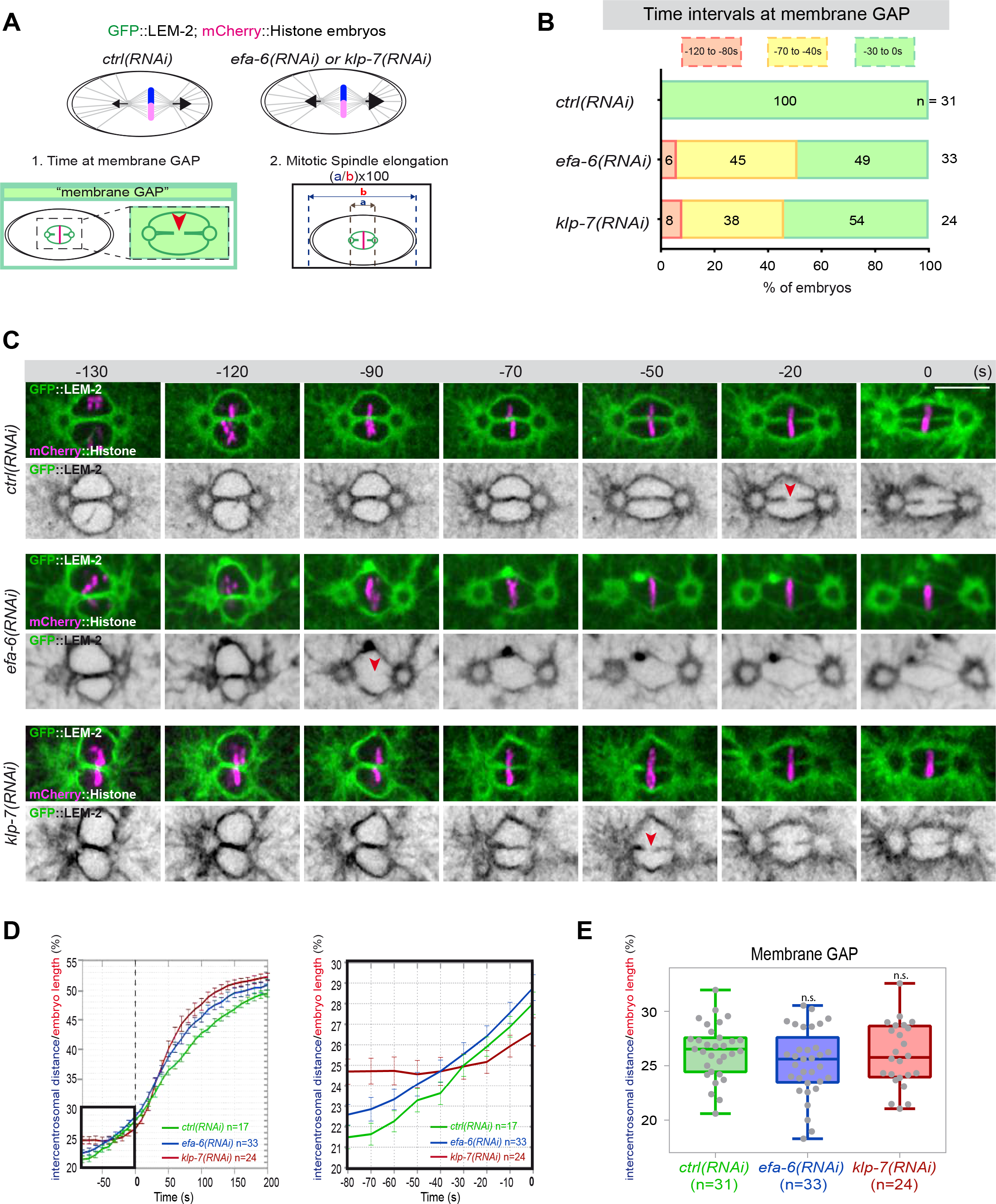
Premature pronuclear membranes scission event upon excessive astral microtubule pulling forces. **A-** Schematics of the approach to test the effect of excessive astral microtubule pulling forces on pronuclear membranes scission (“membrane gap”) (1) and mitotic spindle elongation (2). **B-** Percentage of embryos presenting the pronuclear membranes scission event at different time intervals relative to anaphase onset (0s). The number of embryos (n) analyzed is indicated on the graph and was collected from three independent experiments. **C-** Spinning disk confocal micrographs of early embryos expressing mCherry::Histone, GFP::LEM-2 exposed to mock RNAi (*ctrl*), *efa-6(RNAi)* or *klp- 7(RNAi)*. Times are in seconds relative to anaphase onset (0s). The orange arrowheads indicate pronuclear membranes scission. All panels are at the same magnification Scale Bar, 10 μm. **D-** Intercentrosomal distance normalized to embryo length in percentage during mitosis upon exposure to mock RNAi (*ctrl*), *efa-6* or *klp-7(RNAi)*. Times are in seconds relative to anaphase onset (0s). The graph on the right is a zoom of the first graph focused on the 80 seconds before anaphase onset (0s). **E-** Box and Whisker plots presenting the intercentrosomal distance normalized to embryo length in percentage at the time of pronuclear membranes gap formation in embryos of the indicated genotypes. n= number of embryos analyzed.

### Lamina depolymerization and chromosome alignment are prerequisites for membrane gap formation even in the presence of excessive pulling forces

Embryos with a stabilized lamina, expressing the non-phosphorylable LMN-1 8A, are systematically defective in pronuclear envelopes scission (Velez-Aguilera et al., 2020). We reasoned that the lamina when stabilized, in addition to constituting a physical barrier between chromosomes, could oppose the pulling forces exerted by astral microtubules and could thus prevent elongation of the mitotic spindle during anaphase. To test this model, we measured mitotic spindle elongation in wild-type versus *lmn-1* 8A mutant embryos by measuring the intercentrosomal distance during mitosis. In contrast to wild-type, the spindle did not elongate to the same extent in *lmn-1* 8A mutant embryos, (**Figure 5A, Table 5 Source Data 5**), indicating that stabilization of the lamina interferes with mitotic spindle elongation and thus that lamina depolymerization facilitates spindle elongation.

**Figure 5:**
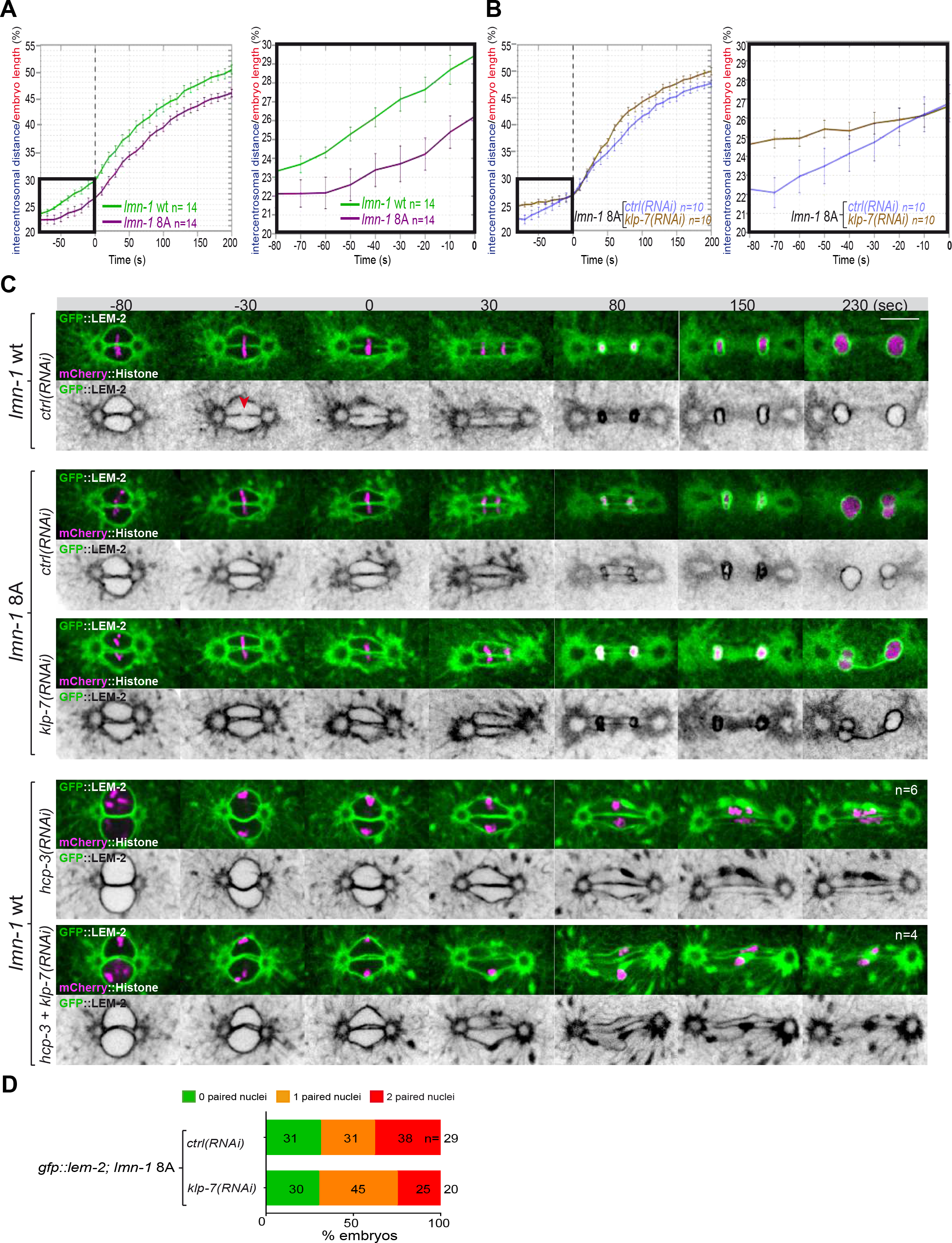
Chromosome alignment and lamina depolymerization are prerequisites for pronuclear membranes scission. **A-** Graphs showing the intercentrosomal distance normalized to embryo length in percentage in wild-type and *lmn-1* 8A embryos during mitosis. Times are in seconds relative to anaphase onset (0s). The graph on the right is a zoom of the first graph focused on the 80 seconds before anaphase onset (0s) **B-** Graphs showing the intercentrosomal distance normalized to embryo length in percentage in *lmn-1* 8A embryos exposed to mock RNAi (*ctrl*) or *klp-7(RNAi)* during mitosis. Times are in seconds relative to anaphase onset (0s). The graph on the right is a zoom of the first graph focused on the 80 seconds before anaphase onset (0s) **C-** Spinning disk confocal micrographs of one-cell stage embryos of the indicated genotype expressing the inner nuclear membrane protein GFP::LEM-2 (shown alone, and in green in the merged images) and mCherry::Histone (magenta, in the merged image). Times are in seconds relative to anaphase onset (0s). The fraction of embryos that showed the presented phenotype is indicated at the bottom right of each image. All panels are at the same magnification Scale Bar, 10 μm. The orange arrowheads point to the pronuclear envelopes scission event. **D-** Percentage of embryos presenting 0 (green bars), 1 (orange bars) or 2 (red bars) paired nuclei at the two-cell stage upon exposure to mock RNAi (*ctrl*) or *klp-7(RNAi).* The number of embryos analyzed (n) is indicated on the graph and was collected from three independent experiments.

We then asked whether excessive microtubule pulling forces might rescue pronuclear envelopes scission in embryos defective in lamina depolymerization. To test this hypothesis, we inactivated *klp-7* in *lmn-1* 8A embryos expressing GFP::LEM- 2 and mCherry::Histone and monitored the nuclear envelope dynamics and mitotic spindle elongation by spinning disk confocal microscopy. While these treatments rescued mitotic spindle elongation defects of *lmn-1* 8A embryos (**Figure 5B, Table 5 Source Data 5**), they failed to restore the formation of pronuclear membranes scission and single nucleus embryos at the 2-cell stage (**Figure 5C, 5D**).

These observations indicate that even premature and excessive spindle elongation is not sufficient to induce membrane scission when the lamina is stabilized during mitosis.

Collectively, these observations indicate that lamina depolymerization is prerequisite for pronuclear envelopes scission.

Previous work has shown that a defect in chromosome alignment on the metaphase plate prevents pronuclear membranes scission (Rahman et al., 2015). In the *efa-6* or *klp-7(RNAi)* embryos with enhanced astral pulling forces, pronuclear membranes scission often occurred before chromosomes were fully aligned on the metaphase plate (**Figure 4C** **and S4C**) suggesting that membrane scission between the pronuclei may not require full and complete chromosome alignment on the metaphase plate.

To further investigate this possibility, we examined whether excessive pulling forces can trigger pronuclear envelopes scission between the pronuclei in the total absence of chromosome alignment. To this end, we inactivated *klp-7* and the essential kinetochore protein CENPA^HCP-3^ to prevent chromosome congression on the metaphase plate (Oegema et al., 2001). In these embryos, no membrane scission was detected despite the extensive mitotic spindle elongation (**Figure 5C**). These observations indicate that chromosome localization in the vicinity of the pronuclei membrane is necessary for pronuclear membranes scission.

### The essential role of PLK-1 in pronuclear envelopes scission is to promote lamina depolymerization

PLK-1 is critically required for pronuclear envelopes scission (Rahman et al., 2015; Martino et al., 2017). We previously showed that LMN-1 is a key PLK-1 target in this process. Consistently, the sole expression of the non-phosphorylable *lmn-1* 8A allele is sufficient to prevent pronuclear envelopes scission, resulting in the formation of embryos with a paired-nuclei phenotype (Velez-Aguilera et al., 2020). Whether PLK-1 regulates pronuclear envelopes scission by targeting other substrates is currently unclear. For instance, PLK-1 could regulate membrane scission by promoting mitotic spindle elongation, or, by activating a factor essential for pronuclear envelopes scission. Recent Focused Ion Beam-Scanning Electron Microscopy (FIB-SEM) analysis has revealed that the four membranes of the pronuclei fuse and become two via a novel membrane structure, the three-way sheet junctions (Rahman et al., 2020). These junctions are absent in *plk-1*ts embryos (Rahman et al., 2020) raising the possibility that PLK-1 could directly regulate their formation.

To discriminate between these hypotheses, we asked whether partial *lmn-1* inactivation by RNAi is sufficient to restore pronuclear envelopes scission in *plk-1*ts embryos (**Figure 6A**). If *lmn-1* inactivation in *plk-1*ts does not restore pronuclear envelopes scission, this would argue that PLK-1 has, beyond lamina depolymerization, other roles to promote membrane gap formation. However, if *lmn-1* inactivation is sufficient to restore membrane gap formation in *plk-1*ts, this would suggest that LMN-1 is possibly the only PLK-1 target in this process, unless partial *lmn-1* inactivation, indirectly affects an inhibitor of pronuclear envelopes scission.

**Figure 6:**
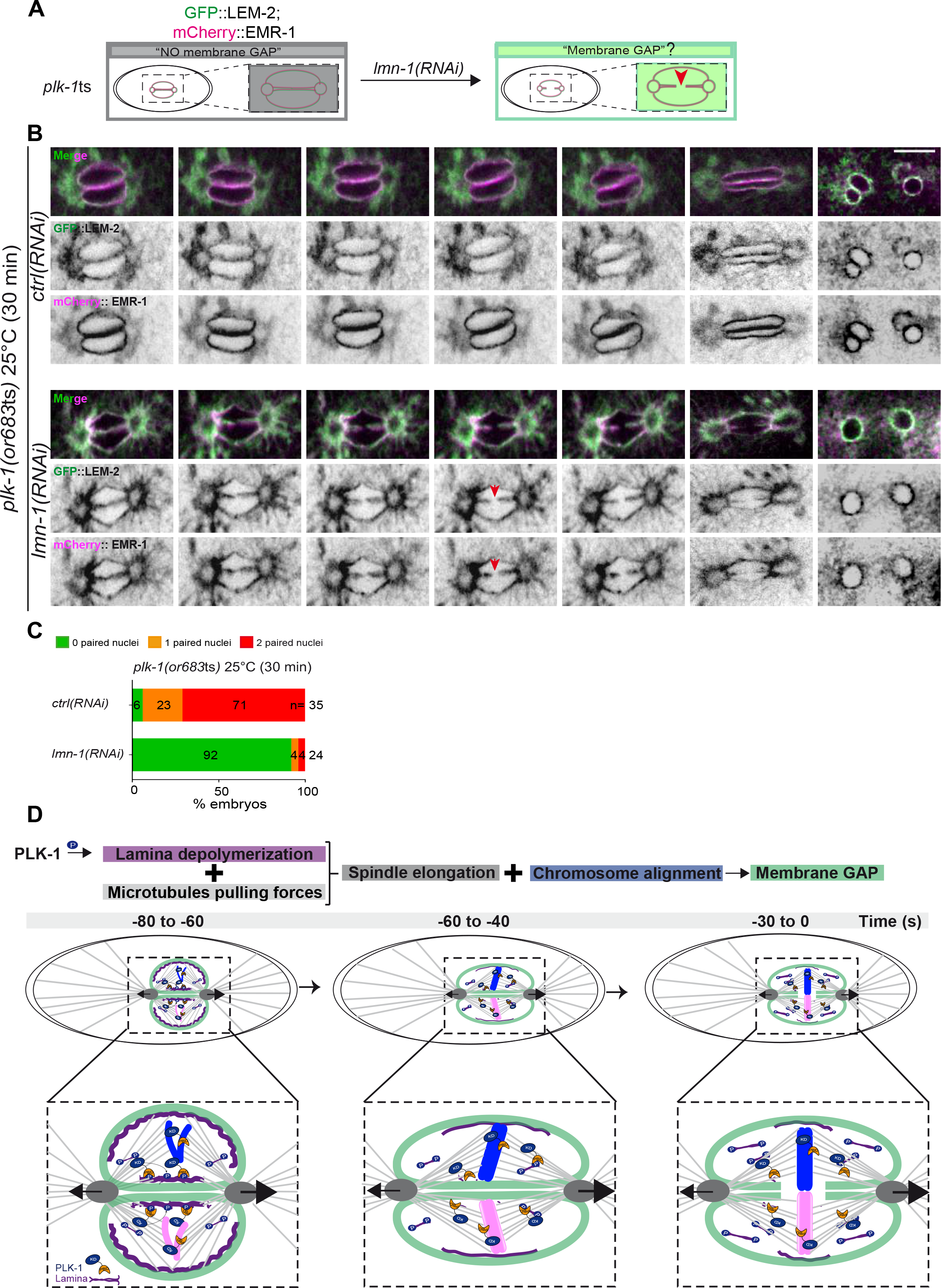
PLK-1 triggers pronuclear membranes scission mainly via lamina depolymerization. **A-** The schematics present the approach used to test whether partial *lmn-1* inactivation in *plk-1*ts embryos expressing GFP::LEM-2 and mCherry::EMR-1 is sufficient to restore membrane gap formation. **B-** Spinning disk confocal micrographs of *plk-1*ts early embryos expressing mCherry::EMR-1 and GFP::LEM-2 exposed mock RNAi (*ctrl*) or *lmn-1(RNAi)*. All panels are at the same magnification. Scale Bar, 10 μm. **C-** Percentage of embryos presenting 0 (green bars), 1 (orange bars) or 2 (red bars) paired nuclei at the two-cell stage upon exposure to mock RNAi (*ctrl*) or *lmn-1(RNAi).* The number of embryos analyzed (n) is indicated on the graph and was collected from three independent experiments. **D-** Working model: Temporal coordination between lamina depolymerization, chromosome alignment, and mitotic spindle elongation is required for pronuclear envelopes scission and parental genomes unification in the early *C. elegans* embryo.

To monitor pronuclear envelopes scission in *plk-1*ts mutant embryos, we constructed a *plk-1*ts strain co-expressing GFP::LEM-2 and mCherry::EMR-1 (EMERIN), which both localize to the inner nuclear envelope and directly interact with the lamina (**Figure 6B**). We then used spinning disk confocal microscopy to monitor pronuclear membranes configuration in one-cell embryos. As reported previously, pronuclear envelopes scission between the pronuclei was totally prevented in *plk-1*ts embryos at restrictive temperature (Rahman et al., 2020; Rahman et al., 2015). However, partial

RNAi-mediated *lmn-1* inactivation was sufficient to restore pronuclear envelopes scission, without affecting the levels and localization of GFP::LEM-2 or mCherry::EMR-1. *lmn-1* inactivation also suppressed the double paired nuclei phenotype of *plk-1*ts embryos in these conditions (**Figure 6C, Table 6 Source Data 6)**, as reported previously (Martino et al., 2017; Rahman et al., 2015). These observations thus suggest that LMN-1 might be the only PLK-1 target involved in regulating pronuclear membranes scission.

In conclusion, our studies indicate that cortical microtubule pulling forces, together with PLK-1, facilitate pronuclear envelopes scission and thereby contribute to the union of the parental chromosomes after fertilization. Our data also clearly indicate that lamina disassembly is a prerequisite for pronuclear membranes scission, even in the presence of excessive cortical microtubule pulling forces that extend the mitotic spindle. Likewise, correct positioning of the parental chromosomes in the vicinity of the pronuclear envelopes is essential for membrane scission. These observations indicate that temporal coordination between lamina depolymerization, chromosome alignment, and mitotic spindle elongation is required for pronuclear envelopes scission in the early *C. elegans* embryo (**Figure 6D**).

## Acknowledgments

We thank P. Moussounda for her help with media preparation. We thank X. Baudin and J. Dumont for microscopy data acquisition and V. Contremoulins for image analysis. We thank R. Karess for critical reading of the manuscript. We acknowledge the ImagoSeine core facility of the Institut Jacques Monod, member of IBiSA and France-BioImaging (ANR-10-INBS-04) infrastructures and the Institut Jacques Monod ‘Structural and Functional proteomic platform”. GVA was supported by the Labex “Who am I?” Laboratory of Excellence No. ANR-11-LABX-0071, the French Government through its Investments for the Future program operated by the French National Research Agency (ANR) under Grant no. ANR-11-IDEX-0005-01 and by the CONACYT grant CVU 364106 and the SECTEI/162/2021 fellowship. NJ is supported by a funding ‘Dynamic Research’ from ANR-18-IDEX-0001, IdEx Université de Paris. Work in the laboratory of LP is supported by grants from “Agence Nationale pour la Recherche” (ANR, France - ANR-17-CE13-0011) and by the “Ligue Nationale Contre le Cancer” (Equipe labéllisée, France).

## Tables Source Data

**Table 1_Source Data 1 related to** **Figure 1**. Quantification of: **1D.** The steps of pronuclear envelopes breakdown: pronuclei permeabilization, lamina disassembly (at the poles, between pronuclei), chromosomes alignment and pronuclear envelopes scission event (membrane gap). **1E.** The surface area occupied by the pronuclei starting 200s before anaphase onset (0s). **1F.** The intercentrosomal distance normalized to embryo length starting 80s before anaphase onset (0s).

**Table 2_Source Data 2 related to** **Figure 2**. Quantification of: **2A.** Percentage of *lmn-1Δ* ; *gfp::lmn-1* and *lmn-1Δ* ; *gfp::lmn-1 8A* embryos presenting 0, 1 or 2 paired nuclei at the two-cell stage upon exposure to mock RNAi (*ctrl*) or *gpr-1/2(RNAi)*. **2E.** GFP::LMN-1 or GFP::LMN-1 8A (**2F-G**) signal intensity during mitosis upon exposure to mock RNAi (*ctrl*), *gpr-1/2(RNAi)* or *klp-7(RNAi)*.

**Table 3_Source Data 3 related to** **Figure 3**. Quantification of: **3C.** Percentage of embryos presenting a pronuclear envelopes scission event **3D.** The intercentrosomal distance normalized to embryo length starting 80s before anaphase onset (0s). **3E.** Percentage of *gfp-lem-2* embryos presenting 0, 1 or 2 paired nuclei at the two-cell stage upon exposure to mock RNAi (*ctrl*) or *gpr-1/2(RNAi)*.

**Table 4_Source Data 4 related to** **Figure 4**. Quantification of: **4B.** Percentage of embryos presenting a pronuclear membranes scission event at different time intervals relative to anaphase onset (0s). **4D.** The intercentrosomal distance normalized to embryo length starting 80s before anaphase onset (0s). **4E.** The intercentrosomal distance normalized to embryo length in percentage at the time of pronuclear membranes gap formation in embryos upon exposure to mock RNAi (*ctrl*), *efa-6(RNAi)* or *klp-7(RNAi)*.

**Table 5_Source Data 5 related to** **Figure 5**. Quantification of: **5A.** The intercentrosomal distance normalized to embryo length starting 80s before anaphase onset (0s) in *lmn-1* wt and *lmn-1* 8A embryos. **5B.** The intercentrosomal distance normalized to embryo length starting 80s before anaphase onset (0s) in *lmn-1* 8A embryos upon exposure to mock (control, ctrl) or *klp-7(RNAi)*. **5E.** Percentage of *gfp- lem-2; lmn-1 8A* embryos presenting 0, 1 or 2 paired nuclei at the two-cell stage upon exposure to mock RNAi (*ctrl*) or *gpr-1/2(RNAi)*.

**Table 6_Source data 6 related to** **Figure 6**. Quantification of **6C.** Percentage of *plk- 1(or683ts)* embryos, expressing GFP::LEM-2 and mCherry::EMR-1 presenting 0, 1 or 2 paired nuclei at the two-cell stage upon exposure to mock RNAi (*ctrl*) or *lmn- 1(RNAi)*.

## Material and Methods

### CONTACT FOR REAGENT AND RESOURCE SHARING

Further information and requests for reagents may be directed to and will be fulfilled by the lead contact author L. Pintard

### EXPERIMENTAL MODEL AND SUBJECT DETAILS

*C. elegans* and bacterial strains used in this study are listed in the Key Resources Table.

#### METHOD DETAILS

##### Molecular Biology

The plasmids and oligonucleotides used in this study are listed in the Key resource table. Gateway cloning was performed according to the manufacturer’s instructions (Invitrogen). All the constructs were verified by DNA sequencing (GATC-Biotech).

##### Nematode strains and RNAi

*C. elegans* strains were cultured and maintained using standard procedures (Brenner, 1974). NPP-22 N-terminally tagged with mCherry using CRIPR/Cas9 was generated by SunyBiotech. The strain expressing LMN-1 N-terminally tagged with mCherry was constructed by mos1-mediated single copy insertion (mosSCI) (Frokjaer-Jensen et al., 2008). The engineered *lmn-1* gene contains a reencoded region in exon 4 essentially as described (Penfield et al., 2018).

RNAi was performed by the feeding method using HT115 bacteria essentially as described (Kamath et al., 2001), except that 2 mM of IPTG was added to the NGM plates and in the bacterial culture just prior seeding the bacteria. As a control, animals were exposed to HT115 bacteria harboring the empty feeding vector L4440 (mock RNAi). RNAi clones were obtained from the Arhinger library (Open Source BioScience) or were constructed.

Feeding RNAi was performed as follows:

Mild *gpr-1/2(RNAi)* was obtained by feeding L4 animals 14-16h at 20°C, whereas strong *gpr-1/2(RNAi)* was obtained by feeding L1 animals for 72 hours at 20°C before filming embryos.

For *efa-6* and *klp-7* inactivation, L4 animals were fed with bacteria at 15°C and embryos were filmed at 23°C.

For single *hcp-3* and *klp-7* inactivation or double *hcp-3*/*klp-7* inactivation, L4 larvae were fed for 14-16h at 15°C with bacteria producing *hcp-3* or *klp-7* dsRNA mixed volume to volume with control bacteria, or with bacteria producing *hcp-3* and *klp-7* dsRNA mixed volume to volume respectively. The embryos were filmed at 23°C.

*plk-1*ts animals were fed with bacteria producing *lmn-1* dsRNA at 15°C from the L1 stage and briefly shifted at 25°C for 30 min before filming the embryos.

##### Microscopy

For the analysis of the paired nuclei phenotype in live specimens by differential interference contrast (DIC) microscopy, embryos were obtained by cutting open young adult hermaphrodites using two 21-gauge needles. Embryos were handled individually and mounted on a coverslip in 3 μl of M9 buffer. The coverslip was placed on a 3% agarose pad. DIC images were acquired by an Axiocam Hamamatsu ICc 1 camera (Hamamatsu Photonics, Bridgewater, NJ) mounted on a Zeiss AxioImager A1 microscope equipped with a Plan Neofluar 100×/1.3 NA objective (Carl Zeiss AG, Jena, Germany), and the acquisition system was controlled by Axiovision software (Carl Zeiss AG, Jena, Germany). Images were acquired at 10- sec intervals.

Live imaging was performed at 23°C using a spinning disk confocal head (CSU-X1; Yokogawa Corporation of America) mounted on an Axio Observer.Z1 inverted microscope (Zeiss) equipped with 491- and 561-nm lasers (OXXIUS 488 nm 150 mW, OXXIUS Laser 561 nm 150 mW) and sCMOS PRIME 95 camera (Photometrics). Acquisition parameters were controlled by MetaMorph software (Molecular Devices). In all cases a 63×, Plan-Apochromat 63×/1.4 Oil (Zeiss) lens was used. Images were acquired at 2-sec or 10-sec intervals. Captured images were processed using ImageJ and Photoshop.

##### Quantification and statistical analysis

GFP::LMN-1 and GFP::LMN-1 8A intensity over time was measured using the Image J software in control and RNAi conditions after background subtraction. Anaphase onset was defined as time 0. Data points on the graphs are the mean of the normalized GFP intensity measurements in control and RNAi conditions for the same define region of interest (ROI). To allow direct comparison between control and RNAi conditions, the average signal intensity of GFP::LMN-1 or GFP::LMN-1 8A at the NE 120 s before anaphase was arbitrarily defined as 1.

The pronuclei area was measured using Image J software by thresholding the region of interest bounded by the mCherry::LMN-1 protein and measuring the total area at each time point.

The intercentrosomal distances and embryos lengths were measured using the IMARIS software by manual tracking of the position of the centrosomes and the poles of the embryos, over time. Then we used the following equation to obtain the corresponding intercentrosomal distance and embryo length at each time point:

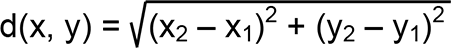

where the first point (first centrosome or embryo pole) is represented by (x_1_,y_1_), and the second point (second centrosome or embryo pole) is represented by (x_2_,y_2_). The data obtained were used to graph the ratio between intercentrosomal distance and embryo length (%) to monitor mitotic spindle elongation.

The results are presented as means ± SEM. The data presented on the graph Figure 4E were compared by Mann-Whitney test. All calculations were performed using GraphPad Prism version 6.00 for Mac OS X, GraphPad Software, La Jolla California USA, www.graphpad.com.

### KEY RESOURCES TABLE

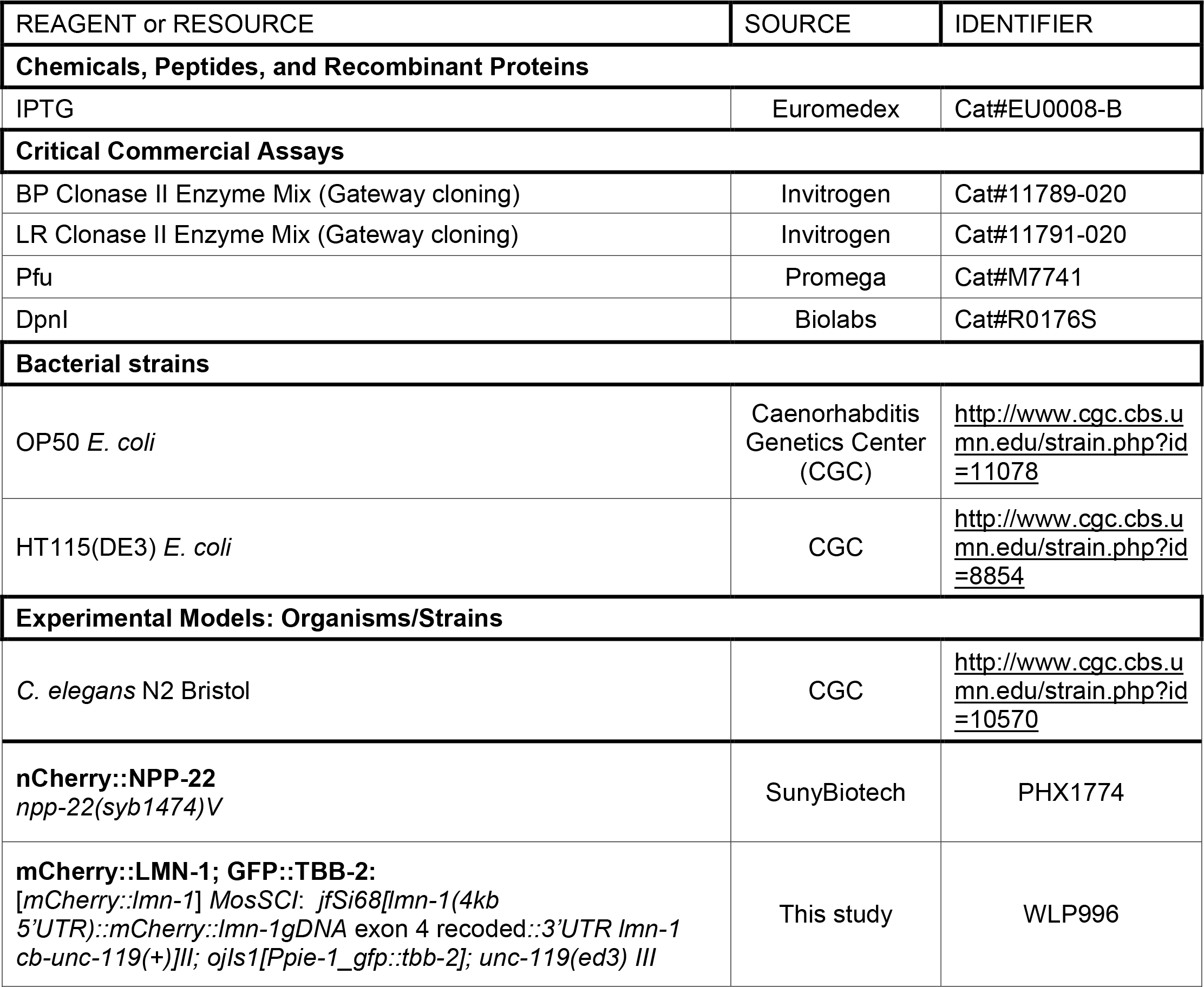
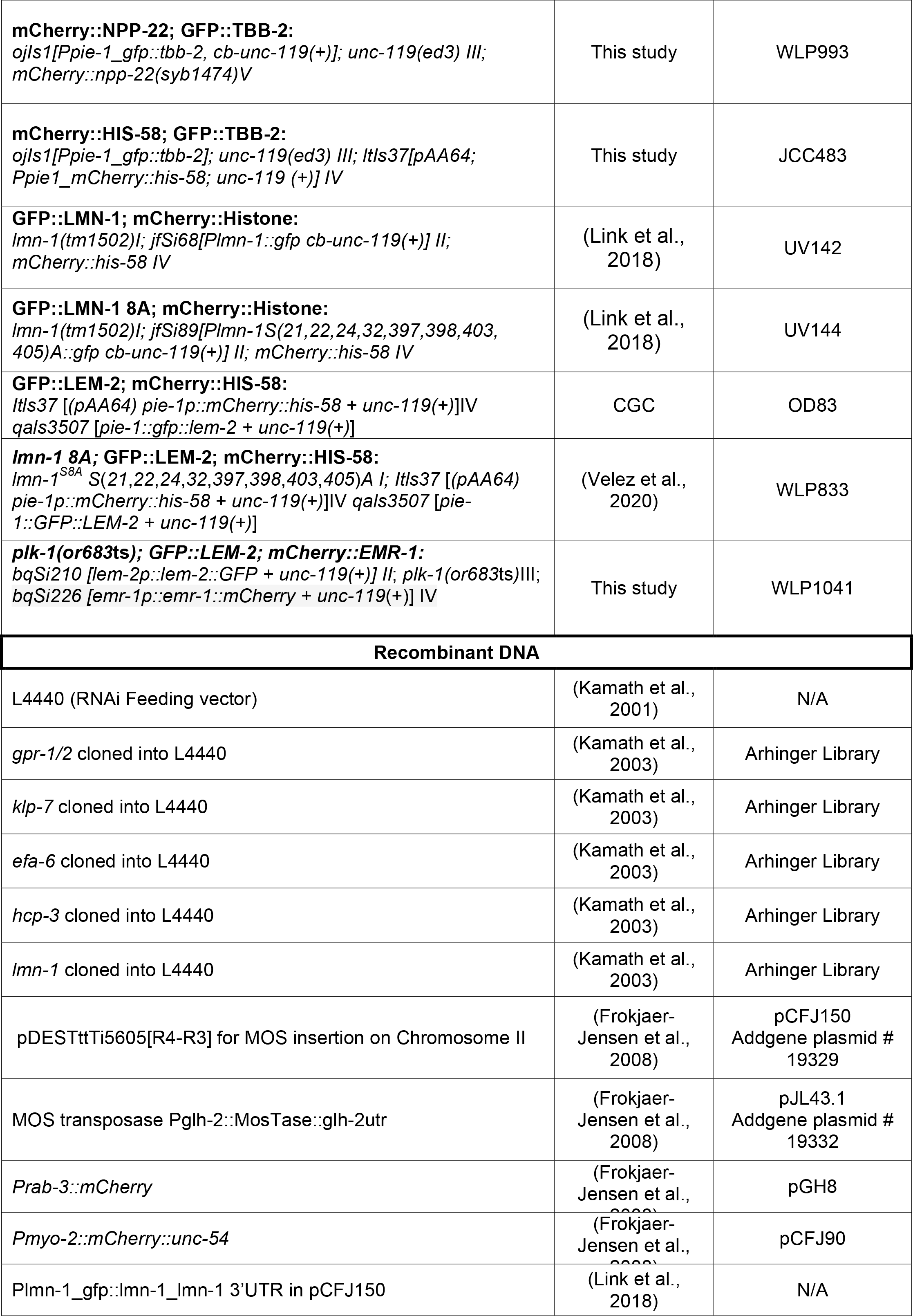
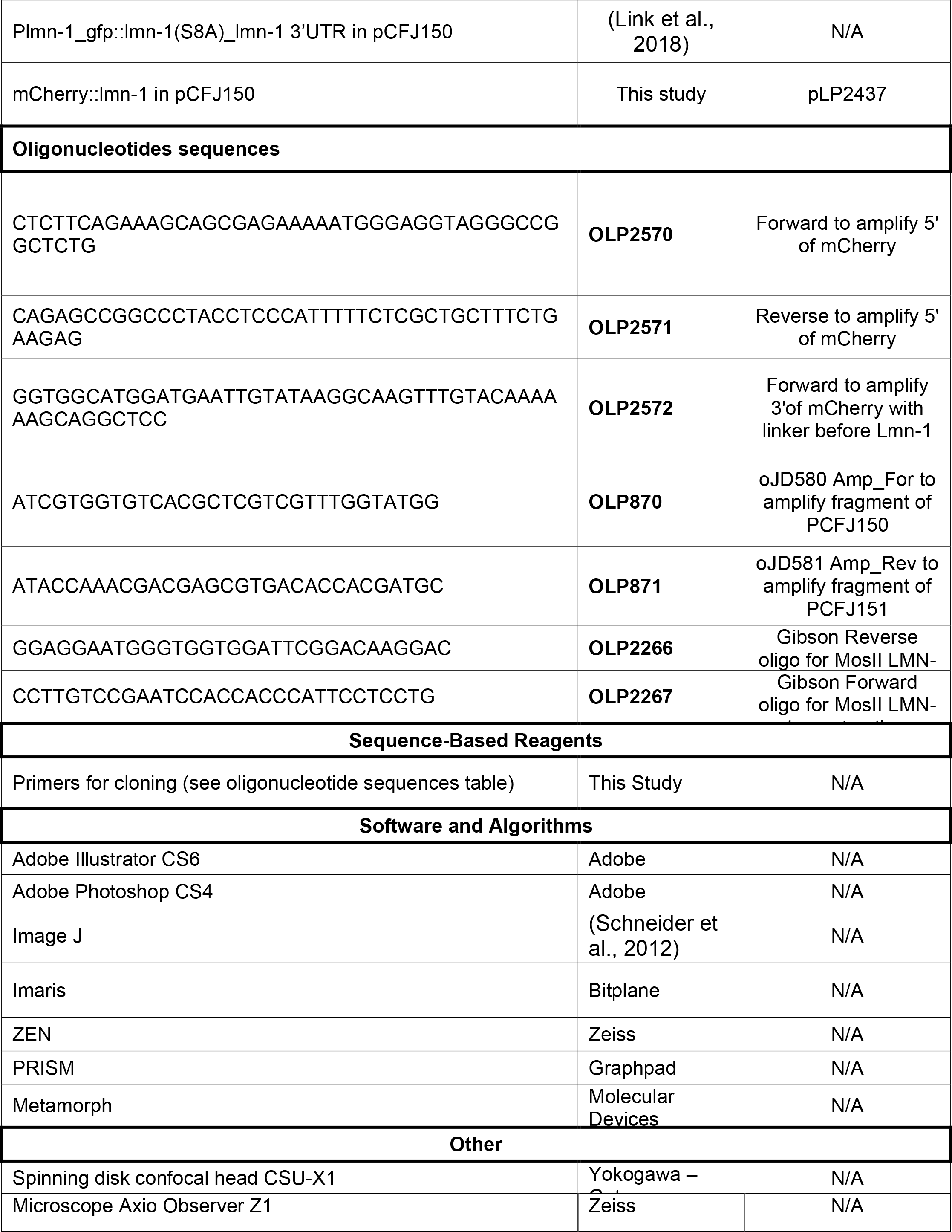

---

**Figure S1:**
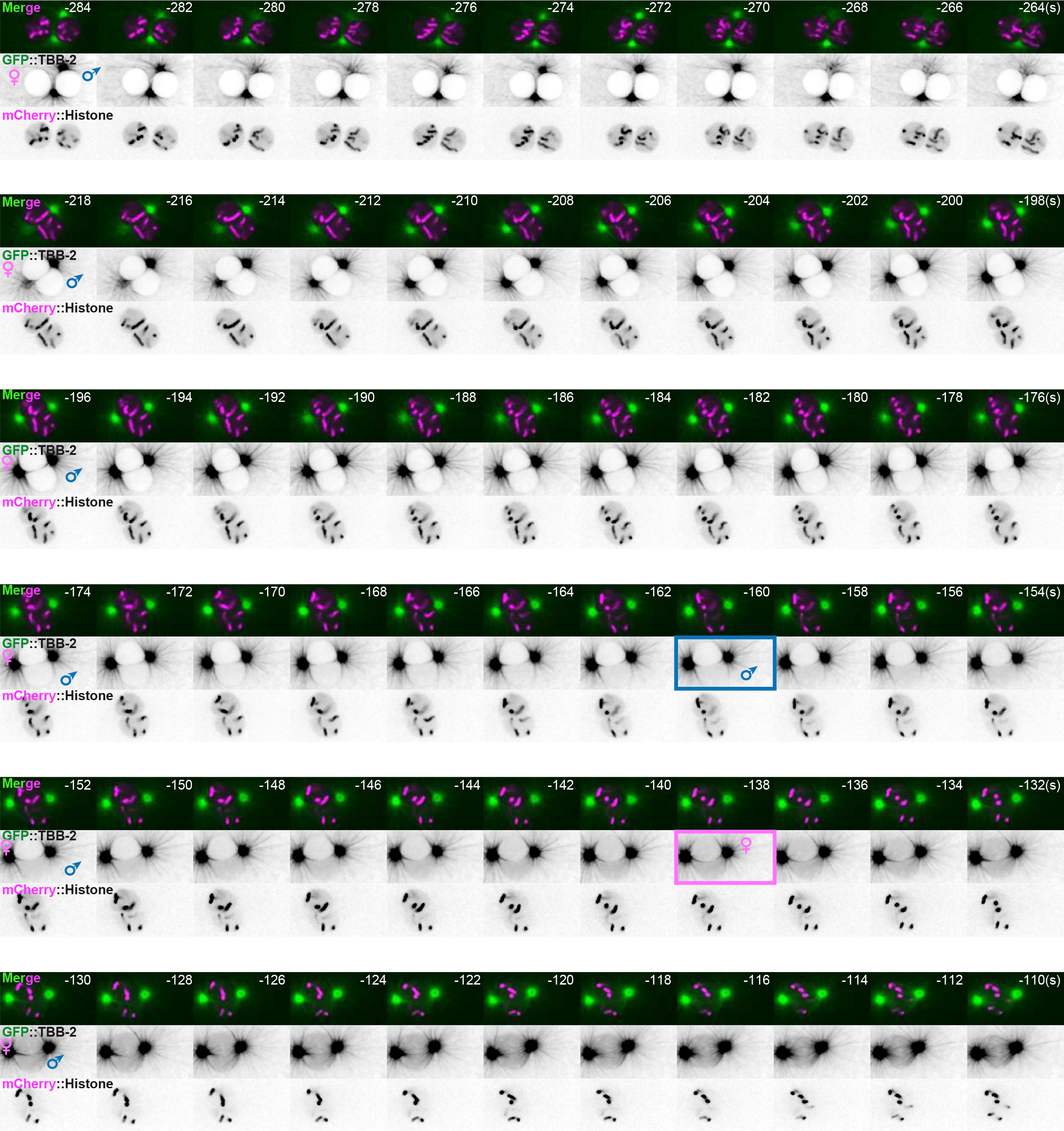
Timing of pronuclei permeabilization visualized by the appearance of soluble tubulin in the pronuclei. Related to figure 1. Spinning disk confocal micrographs of embryos expressing GFP::TBB-2 (shown alone, and in green in the merged images) and mCherry::Histone, (magenta, in the merged image) from pronuclei meeting to membrane permeabilization. The timing of paternal and maternal pronuclei permeabilization is indicated by a blue and pink rectangle respectively. Time interval every 2 seconds, relative to anaphase onset (0s). All panels are at the same magnification. Scale Bar, 10 μm.

**Figure S2:**
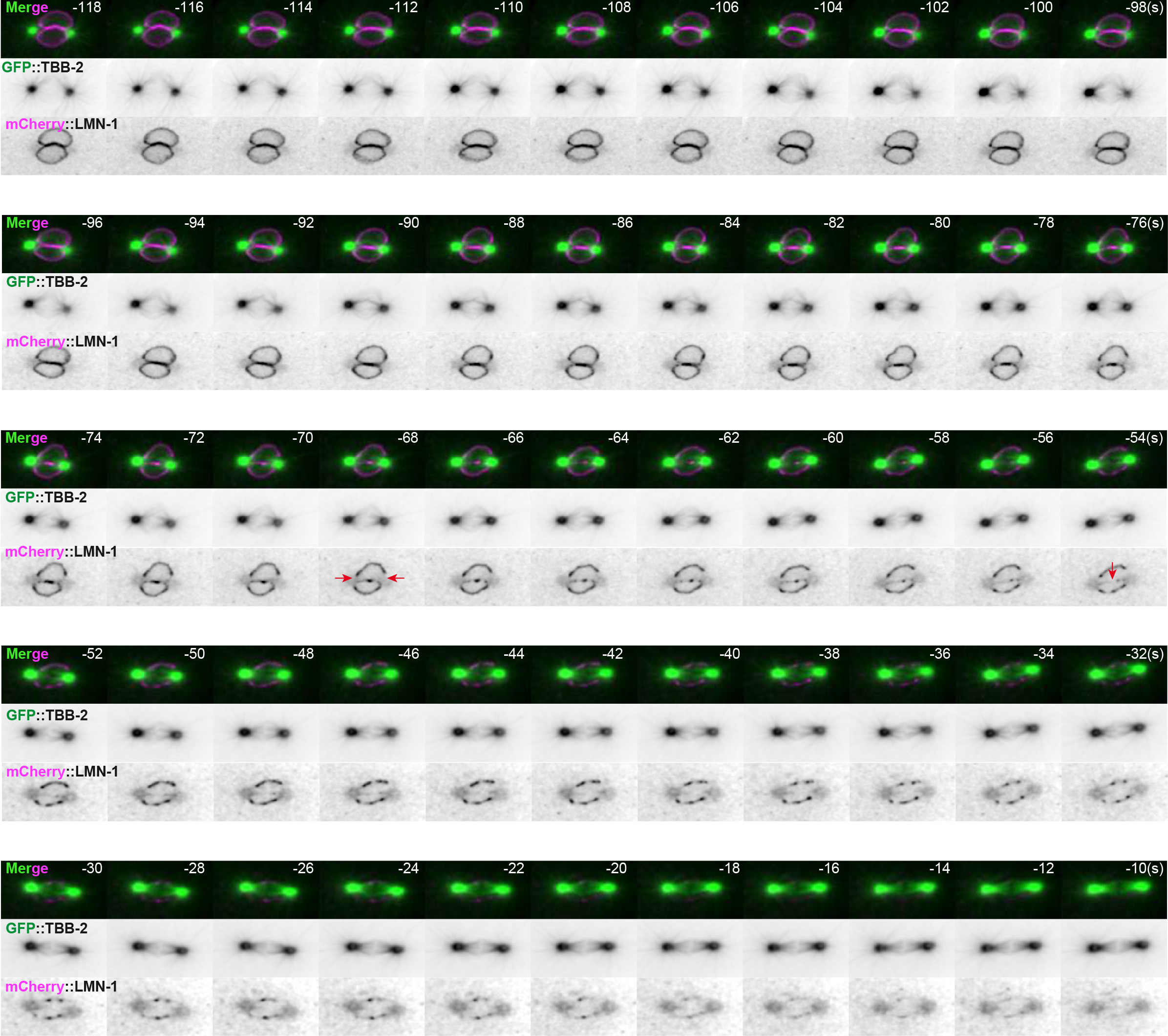
Spatio-temporal lamina depolymerization during mitosis. Related to Figure 1. Spinning disk confocal micrographs of embryos expressing GFP::TBB-2 (shown alone, and in green in the merged images) and mCherry::LMN-1 (shown alone and in magenta in the merged image). Time interval every 2 seconds, relative to anaphase onset (0s). All panels are at the same magnification. Scale Bar, 10 μm.

**Figure S3:**
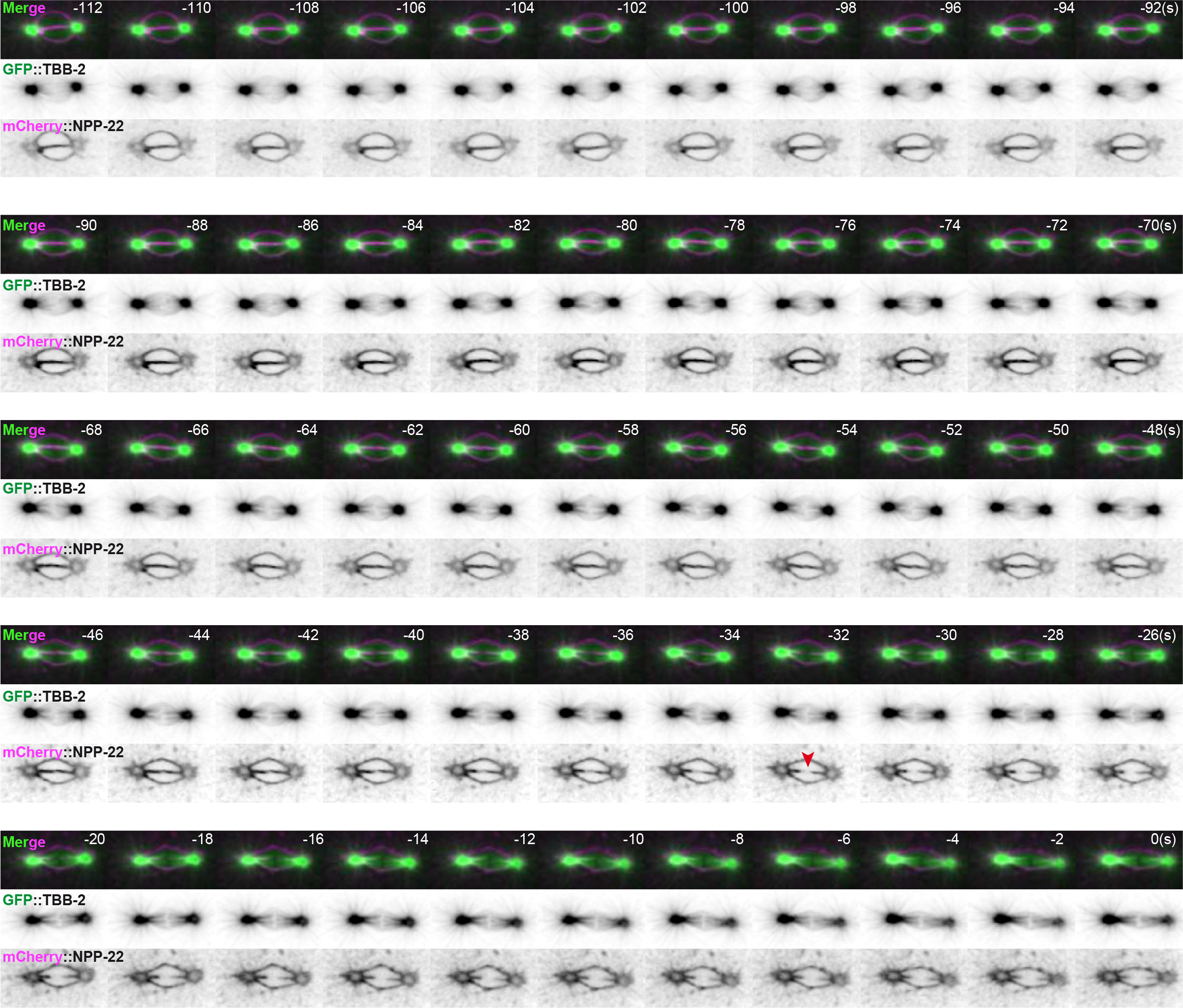
Timing of pronuclear envelopes scission. Related to Figure 1. Spinning disk confocal micrographs of embryos expressing GFP::TBB-2 (shown alone, and in green in the merged images) and mCherry::NPP-22 (shown alone and in magenta in the merged image). Time interval every 2 seconds, relative to anaphase onset (0s). All panels are at the same magnification. Scale Bar, 10 μm.

**Figure S4:**
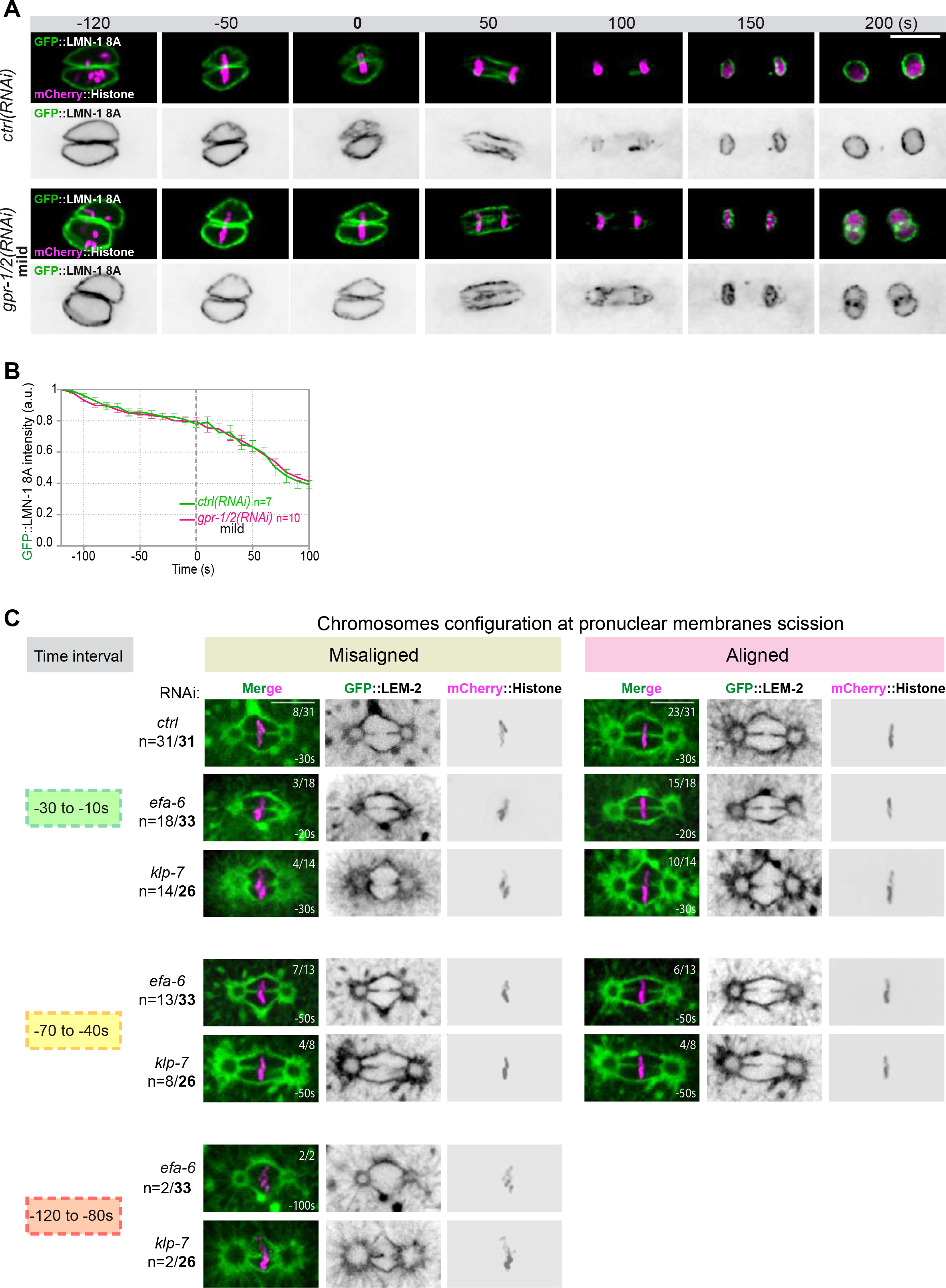
Quantification of GFP::LMN-1 8A and chromosome configuration at the time of membrane gap formation upon an excessive astral microtubule pulling forces. Related to figures 2 and 4. **A-**Spinning disk confocal micrographs of early *lmn-1Δ* mutant embryos expressing GFP::LMN-1 8A (shown alone, and in green in the merged images) and mCherry::Histone (magenta, in the merged image) exposed to mock RNAi (*ctrl*) in the upper panels or mild *gpr1/2(RNAi)* in the lower panels. Times are in seconds relative to anaphase onset (0s). All panels are at the same magnification Scale Bar, 10 μm. **B-** Quantification of GFP::LMN-1 8A signal intensity above background at the nuclear envelope in embryos exposed to mock RNAi (*ctrl*) or mild *gpr1/2(RNAi)* during mitosis. The mean +/- SEM is presented for n embryos of the indicated genotypes. Data were collected from three independent experiments. **C-** Representative spinning disk confocal micrographs of one-cell stage embryos expressing GFP::LEM-2 (shown alone, and in green in the merged images) and mCherry::Histone (magenta, in the merged image) exposed to control, *efa-6* or *klp-7* RNAi. Embryos presenting pronuclear envelopes scission at each time interval are presented along with the configuration of their chromosomes (misaligned or aligned). n= number of embryos presenting a membrane scission at each time interval over the total number of embryos analyzed (in bold). The fraction of embryos presenting aligned or misaligned chromosomes at the time of pronuclear membranes scission is indicated at the top right of each image. Scale Bar, 10 μm.

## Notes

### Competing Interest Statement

The authors have declared no competing interest.

